# Structural and functional insights in flavivirus NS5 proteins gained by the structure of Ntaya virus polymerase and methyltransferase

**DOI:** 10.1101/2023.05.03.539197

**Authors:** Kateřina Krejčová, Petra Krafcikova, Martin Klima, Dominika Chalupska, Karel Chalupsky, Eva Zilecka, Evzen Boura

## Abstract

Flaviviruses are single-stranded positive-sense RNA (+RNA) viruses that are responsible for several (re)emerging diseases such as Yellow, Dengue or West Nile fevers. The Zika epidemic highlighted their dangerousness when a relatively benign virus known since the 1950s turned into a deadly pathogen. The central protein for their replication is NS5 (non-structural protein 5), which is composed of the N-terminal methyltransferase (MTase) domain and the C-terminal RNA-dependent RNA-polymerase (RdRp) domain. It is responsible for both, RNA replication and installation of the 5’ RNA cap. We structurally and biochemically analyzed the Ntaya virus MTase and RdRp domains and we compared their properties to other flaviviral NS5s. The enzymatic centers are well conserved across *Flaviviridae*, suggesting that the development of drugs targeting all flaviviruses is feasible. However, the enzymatic activities of the isolated proteins were significantly different for the MTase domains.

## Introduction

Single-stranded positive-sense RNA (+RNA) viruses are responsible for most of the recent virus outbreaks, local epidemics, and most importantly, the COVID-19 pandemic. Flaviviruses are one of the +RNA virus families that contain relatively benign or animal pathogens as well as dangerous human pathogens. Yellow fever, caused by a flavivirus (Yellow fever virus, YFV) was considered the worst disease of the 19^th^ century and was only contained after a vaccine was developed in the 1930s [1]. Recently, we have witnessed outbreaks of other flaviviruses, most importantly the mosquito-borne West Nile virus (WNV) [2], Dengue virus (DENV) [3] and Zika virus (ZIKV) [4] in the Americas and the tick-borne encephalitis virus (TBEV) in Europe and Asia [5, 6].

Ntaya virus (NTAV) was first isolated from mosquitos in Uganda in 1951 [7]. However, the exact mosquito species that serves as a vector is unknown although the genus *Culex* is the most probable [8]. Together with several other flaviviruses, it comprises the Ntaya virus group, which used to have four other viral species besides NTAV: Bagaza virus (BAGV), Israel turkey meningoencephalitis virus (ITV), Ilheus virus (ILHV), and Tembusu virus (TMUV) [9]. However, recently it was shown that BAGV and ITV are actually the same virus [10]. Antibodies against Ntaya virus have been discovered in a variety of migratory birds [11] and domestic mammals, such as sheep, cattle, goats and pigs [12]. In birds, the virus is neurotropic and causes hemorrhages in the brain and other organs [13]. Antibodies against Ntaya virus have also been discovered in humans form West, Central and East African regions and the virus is suspected to cause an illness that manifests itself with fever and headache [14].

Ntaya virus and other flaviviruses encode several non-structural proteins (NS1, NS2A, NS2B, NS3, NS4A, NS4B and NS5) that ensure their replication in infected cells [15]. Some of them are enzymes: for example NS2B-NS3 is a protease, NS3 is also a helicase and the NS5 protein bears the most important enzymatic activity for an RNA virus - the RNA-dependent RNA-polymerase (RdRp). In addition, the NS5 protein has an N-terminal methyltransferase (MTase) domain that is responsible for RNA cap formation, a process necessary for efficient viral RNA (vRNA) translation and immune evasion [16, 17]. However, the functions of flaviviral NS5 proteins are not fully understood, and only a handful of crystal structures of the RdRp domain are available from the most medically important flaviviruses including Zika, Dengue, West Nile, Japanese encephalitis and Yellow fever viruses [18-22]. The MTase domains are more explored, and crystal structures of MTases from less-known flavivirus such as the Langat virus are available [23]. We aimed to better understand the NS5 protein function. We chose the Ntaya virus NS5 protein for analysis and solved the crystal structures of the RdRp and MTase domains. We also performed a structural and functional comparison of flaviviral RdRps and MTases, which revealed their common features and surprising differences in the enzymatic activities of the MTase domains.

## Results

### Crystal structure of Ntaya RdRp

We aimed to solve the crystal structure of the Ntaya polymerase to gain more insights into the replication of flaviviruses. Eventually we obtained crystals that belonged to the monoclinic P2_1_ spacegroup and diffracted to 2.8Å resolution. The structure was solved by molecular replacement and revealed a fold resembling a cupped human right hand with fingers, palm and thumb, which is typical for viral polymerases (Figure 1). It is a predominantly α-helical fold composed of twenty-seven helices (helices α10 - α36, helices α1 - α9 of the NS5 protein are located in the N-terminal MTase domain) with five small β-sheets. Interestingly, all eleven β-strands (β10 - β20) forming these β-sheets are oriented in an antiparallel manner (Figure 1D). The flaviviral RdRp domain also contains two zinc fingers that are important for the overall fold stability [22]; one is located in the vicinity of helices α10, α14, α16 and α22 and is formed by two cysteine residues (Cys449 and Cys452), one histidine (His444) and one glutamate (Glu440) residues (Cys_2_HisGlu, Figure 1C). This is somewhat different from the canonical Cys_2_His_2_ zinc finger that is widespread in DNA binding motifs [24] However, Glu440 is absolutely conserved among flaviviral RdRps (SI Figure 1). The second zinc finger is localized above the β18-β19 sheet and between helices α33 and α35 and it is formed by cysteine residues Cys733 and Cys852 and histidine residues His717 and His719 (Cys_2_His_2_-type, Figure 1C). The conserved motives A - G that bear most of the catalytically important residues are arranged along the template entry channel (F and G), the active site (A, B, D and E), and the dsRNA exit channel (C), as expected based on their conserved functions: *i*) template binding (B and C), *ii*) incoming nucleotide binding and its stabilization in a proper conformation (E, F, G), *iii*) priming (D) and *iv*) the formation of the phosphodiester bond (A).

**Figure 1:**
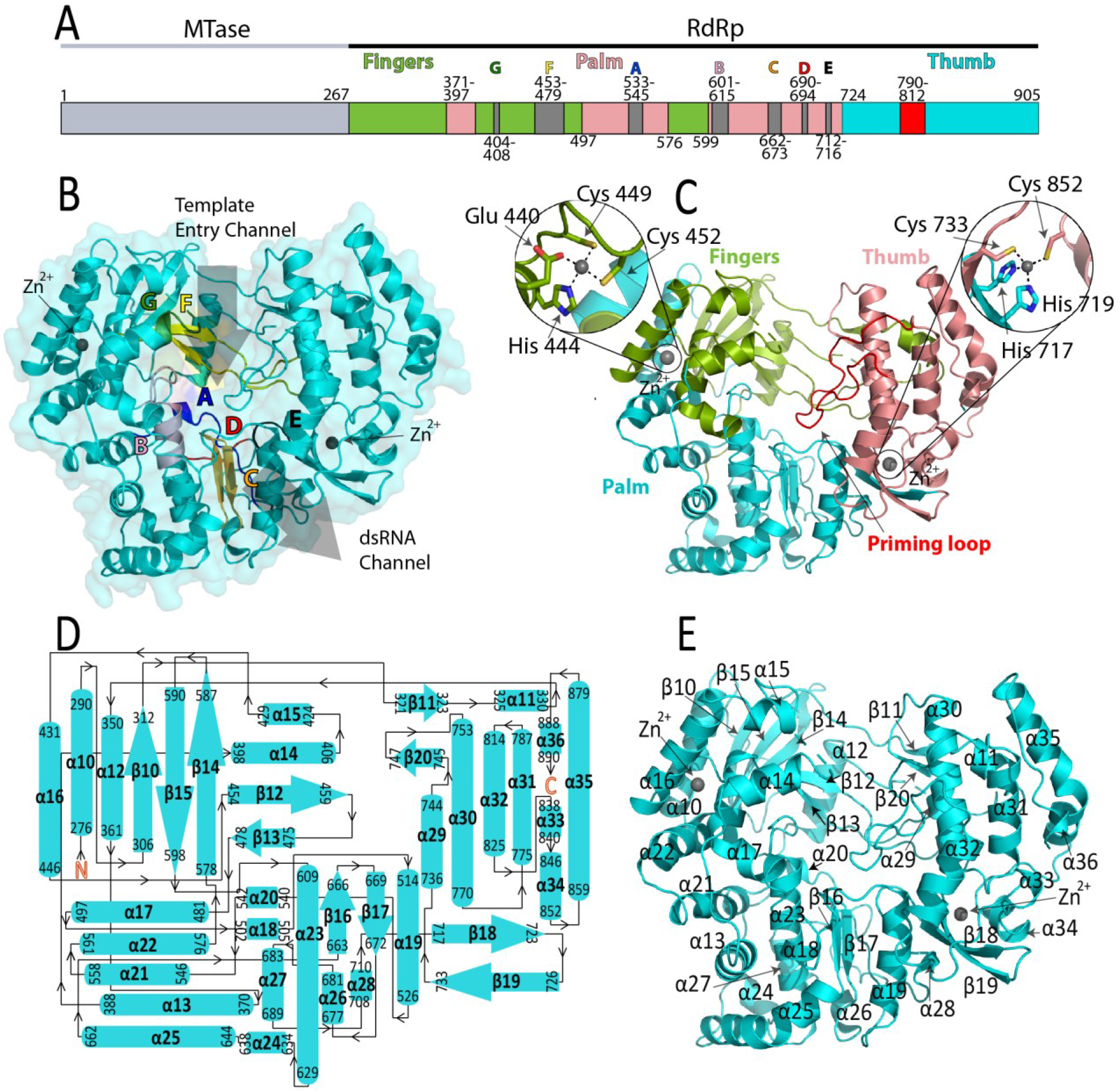
Crystal structure of the Ntaya RdRp domain. A) Schematic representation of the NS5 protein which is composed of the MTase and RdRp domains. Palm, thumb and finger subdomains and motives A-G are shown. B) Overall structure of Ntaya RdRp, template entry and dsRNA channels and motives A-G are highlighted. C) The three subdomains are depicted in different colors: fingers (green), palm (cyan), thumb (pink) and the priming loop (red). Two zinc-binding fingers are zoomed. D) Topological representation of the secondary structure of the Ntaya RdRp. E) Secondary structure elements are labeled.

The overall fold of the Ntaya RdRp domain is in good agreement with previously described flaviviral RdRps (Figure 2A, SI Figure 2). The most similar seems to be the RdRp from the Zika virus (RMSD of superposed structures = 0.866, ΔD_max_ = 1.27Å) while the most different one (RMSD of superposed structures = 1.708, ΔD_max_ = 3.43Å) was the one from the West Nile virus (Figure 2A). Most of the structural differences are in the conformations of loops, among them the priming loop is the most important for the enzymatic function - flaviviral RdRps belong to the primer-independent polymerases. The closed conformation of flaviviral RdRp allows only for the entry of ssRNA and the initiation of RNA synthesis is by the *de novo* mechanism where the priming loop partially fulfils the function of the primer. We examined the conformation of the Ntaya priming loop in detail and compared it to ZIKV and WNV priming loops (Figure 2B, C). While the beginning and end of these priming loops (Trp792 and Glu812) are always in the same conformation the rest significantly differs. The ZIKV priming loop is virtually in the same conformation as Ntaya, the only difference being a different rotamer of Trp800, a residue important for the stabilization of the initiation complex [25]. In contrast, Trp800 is displaced in the case of WNV. Actually, the overall conformation of WNV priming loop is different, another significantly displaced residue is the His803 residue (Figure 2C), which could play a role in stabilizing the initiation complex via a stacking interaction with the base of a priming NTP [25]. Interestingly, the position of Trp808 is absolutely conserved among all analyzed flaviviral polymerases (Figure 2, SI Figure 2) suggesting that this residue is important for the function of the priming loop.

**Figure 2:**
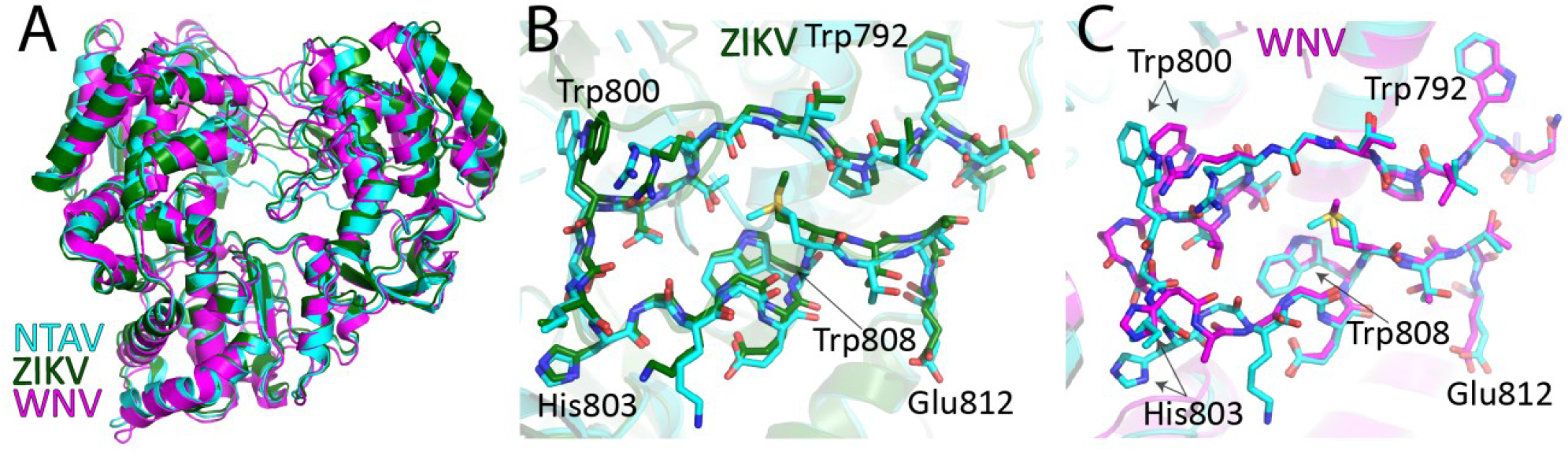
Structural alignment of Flaviviral RdRp domains. A) The overall structural alignment of Ntaya (cyan), Zika (dark green, PDB: 5M2Z) and West Nile (magenta, PDB: 2HFZ) RdRps. B, C) Structural superposition of NTAV (cyan) and ZIKV (dark green) or WNV (magenta) priming loops. The key residue Trp800 (Trp797 in case of ZIKV), which is involved in the stabilization of the priming nucleotide, is indicated.

### Ntaya MTase crystal structure

We also aimed to solve the crystal structure of the MTase domain of NS5. We supplemented the protein with the pan-MTase inhibitor sinefungin and obtained well-diffracting crystals (2.3Å resolution, SI Table 1). The structure was solved by molecular replacement (detailed in the M&M section) and revealed the overall fold of the Ntaya MTase which was in good agreement with previously solved structures of flaviviral MTases [26, 27]. It is a mixed α-β fold (Figure 3B) that resembles a sandwich, where a central β-sheet is surrounded by α-helices (Figure 3). The central β-sheet is composed of seven β-strands (β4, β3, β2, β5, β6, β8 and β7 as viewed from the S-adenosyl-methionine (SAM) binding pocket) and, together with β1 and β9 form β-sheets that resemble the letter *J* (Figure 3C). These *J* β-sheets are well conserved among analysed flaviviral MTases, except for the β3 and β2 strands, where Ntaya has a different conformation (Figure 3C). A three-helix bundle (α1, α2 and α8) contacts and stabilizes the loop connecting β7 and β8 strands, and a four-helix bundle (α6, α5, α4 and α3) together with a small β1 and β9 sheet is located above the central sheet, while helices α7 and α9 are located below.

**Figure 3:**
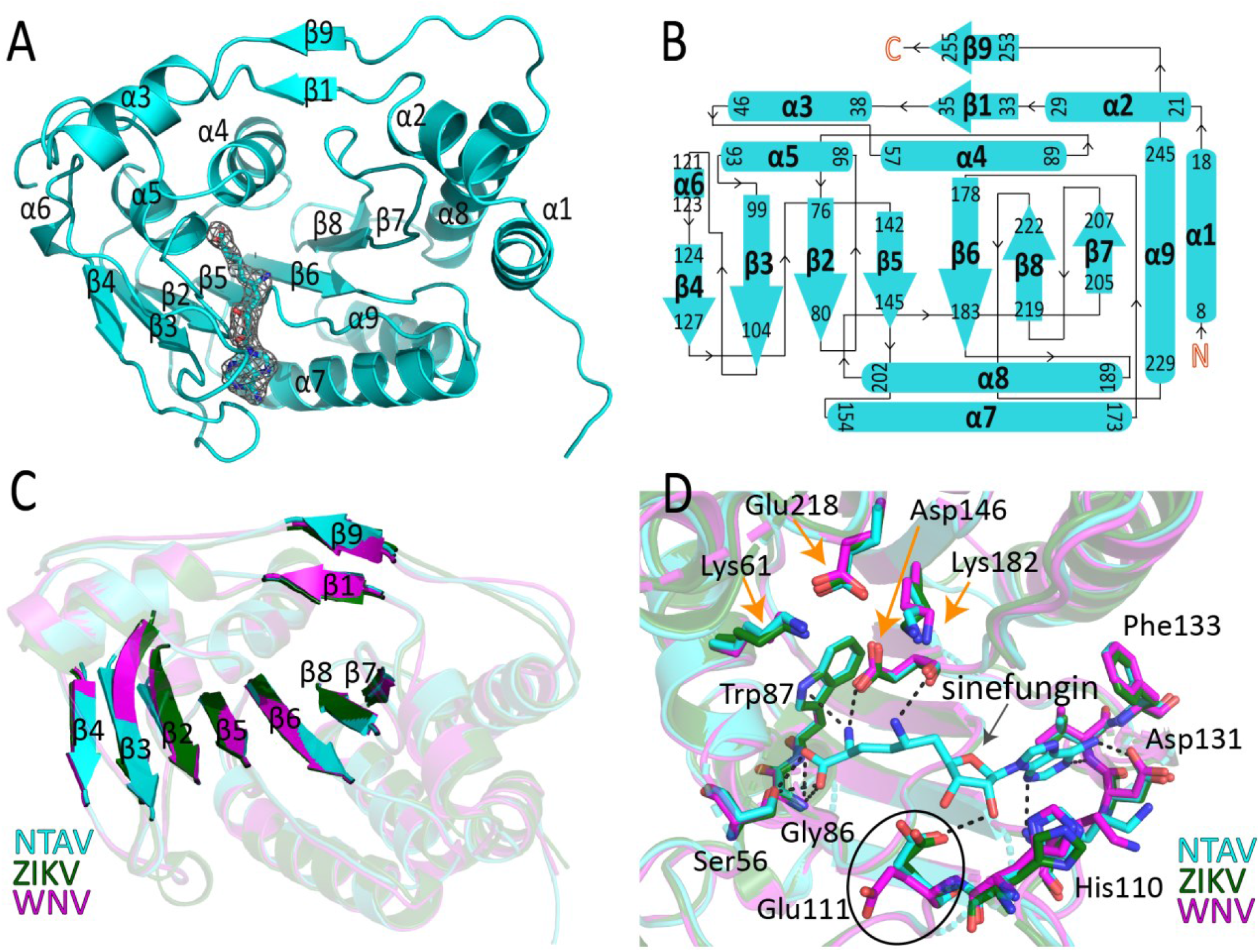
Crystal structure of the Ntaya MTase domain. A) Overall fold of the Ntaya MTase domain with sinefungin bound. An Fo-Fc omit map contoured at 2σ is shown around the sinefungin. B) Topological representation of the Ntaya MTase secondary structure. C) Structural superposition of Ntaya (cyan), Zika (dark green, PDB: 5MRK) and West-Nile (magenta, PDB: 4R8S) MTase domains, the β-sheets are highlighted. D) Structural comparison of the SAM binding pockets. Hydrogen bonds between sinefungin and key residues are shown. Residues of the catalytic tetrad are highlighted by orange arrows.

### Sinefungin binding mode

The electron density for sinefungin was clearly visible upon molecular replacement (Figure 3A). Sinefungin was located in the SAM binding pocket, which is defined by four β-strands (β4, β3, β2 and β5) and three helices (α3, α4, α5). The sinefungin molecule is bound to the SAM binding pocket mainly through hydrogen bonds. The 2′ hydroxyl of the ribose ring forms a hydrogen bond with the side chain of Glu111. The adenosine ring forms hydrogen bonds to the backbones of Lys105 and Val132 and its 6-amino group interacts with the side chain of Asp131. The amino acid moiety of sinefungin is coordinated by hydrogen bonds to Trp87, Asp146, Ser56 and Gly86 (Figure 3D). Superposition of Ntaya and Zika virus MTases revealed that the catalytic tetrad KDKE (residues Lys61, Asp146, Lys182 and Glu218) is in the same conformation (Figure 3D), which is not surprising given the absolute conservation of these residues for all the analyzed flaviviral MTases (SI Figure 1).

### GTP binding mode

The NS5 protein, specifically its MTase domain, is also an RNA guanylyltransferase [28], and thus its MTase domain has a GTP binding site [29]. We were interested in the GTP binding mode and aimed to solve a crystal structure with GTP bound. To begin, crystals of Ntaya MTase were prepared without the presence of sinefungin, resulting in the presence of S-adenosyl-homocysteine (SAH) from bacteria bound in the SAM binding pocket of the recombinant protein. Subsequently, the crystals were soaked overnight with GTP and magnesium as described in the Materials and Methods section. These soaked crystals diffracted at a resolution of 2Å, revealing clear electron density for both ligands (SI Figure 3A), with each ligand localized at its respective site (Figure 4A). The GTP molecule formed hydrogen bonds with key residues within the GTP/cap-binding pocket. The 2′ hydroxyl group of the ribose ring of GTP interacts with the side chains of Gln17 and Lys13. Actually, Lys13 forms hydrogen bonds with both the 2′ and 3′ hydroxyl groups. Also, the main chains of Ser151 and Pro152 are involved in hydrogen bonding with the 3′ hydroxyl group. The 2-amino group of the guanine ring forms hydrogen bond with the backbones of Leu16, Gln17 and Leu19. The phosphate groups are stabilized by hydrogen bonds to Arg28, Arg213 and Ser215 (Figure 4B). The magnesium atom was clearly visible and was coordinated by six oxygen atoms - three from the phosphate groups of GTP (one oxygen from each phosphate group) and three water molecules (SI Figure 3B). In fact, this octahedral coordination is used to distinguish magnesium from water [30]. However, a structural comparison with the crystal structure of Zika MTase bound to GTP revealed a different conformation of the triphosphates (Figure 4C). This is most likely caused by the lack of magnesium in the crystal structure of the Zika MTase/GTP complex in the study of Zhang et al. [31]. Magnesium is present in the cytoplasm where the ZIKV replicates therefore we believe our structure represents the physiological state. We also observed the SAH molecule, and its binding mode was the same as the binding mode of sinefungin with the obvious exception that SAH does not have an amine group that could hydrogen bond with Asp146 (Figure 4D).

**Figure 4:**
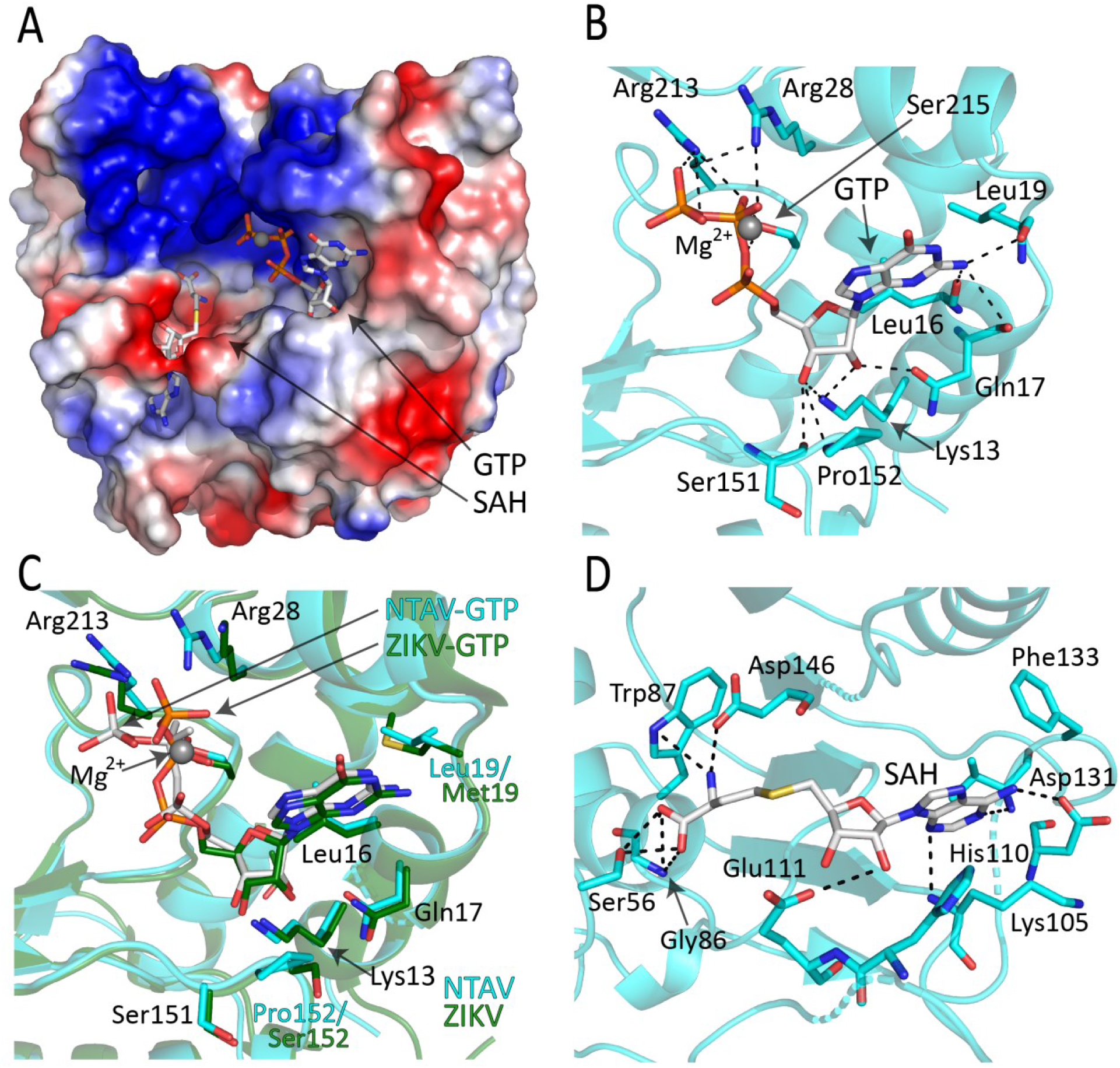
Crystal structure of the Ntaya MTase domain in complex with GTP and SAH. A) SAH and GTP bound to the Ntaya MTase domain. The surface is colored according to the electrostatic potential and SAH and SAM are shown in stick representation. B) A detailed view of GTP bound to key residues of the GTP/cap-binding pocket. Selected hydrogen bonds between GTP and the key residues are depicted and labeled. C) Structural alignment of GTP/cap-binding pocket of Ntaya (cyan) and Zika (dark green, PDB: 5GOZ) with GTP bound. D) A detailed view of SAH bound to key residues of the SAM binding pocket.

### RdRps enzymatic activities

We were also interested in the functional comparison of RdRps from various flaviviruses. We chose the NTAV, JEV, WNV, YF, and ZIKV RdRp domains of NS5 proteins for this comparison. We used a classical primer extension assay, where one primer was fluorescently labeled, and we monitored the progress of the reaction using denaturing PAGE (Figure 5). Consistent with the high structural homology of their active sites, the activity of these enzymes was similar. The most active enzyme was from ZIKV, but all the RdRps exhibited fair activity (Figure 5).

**Figure 5:**
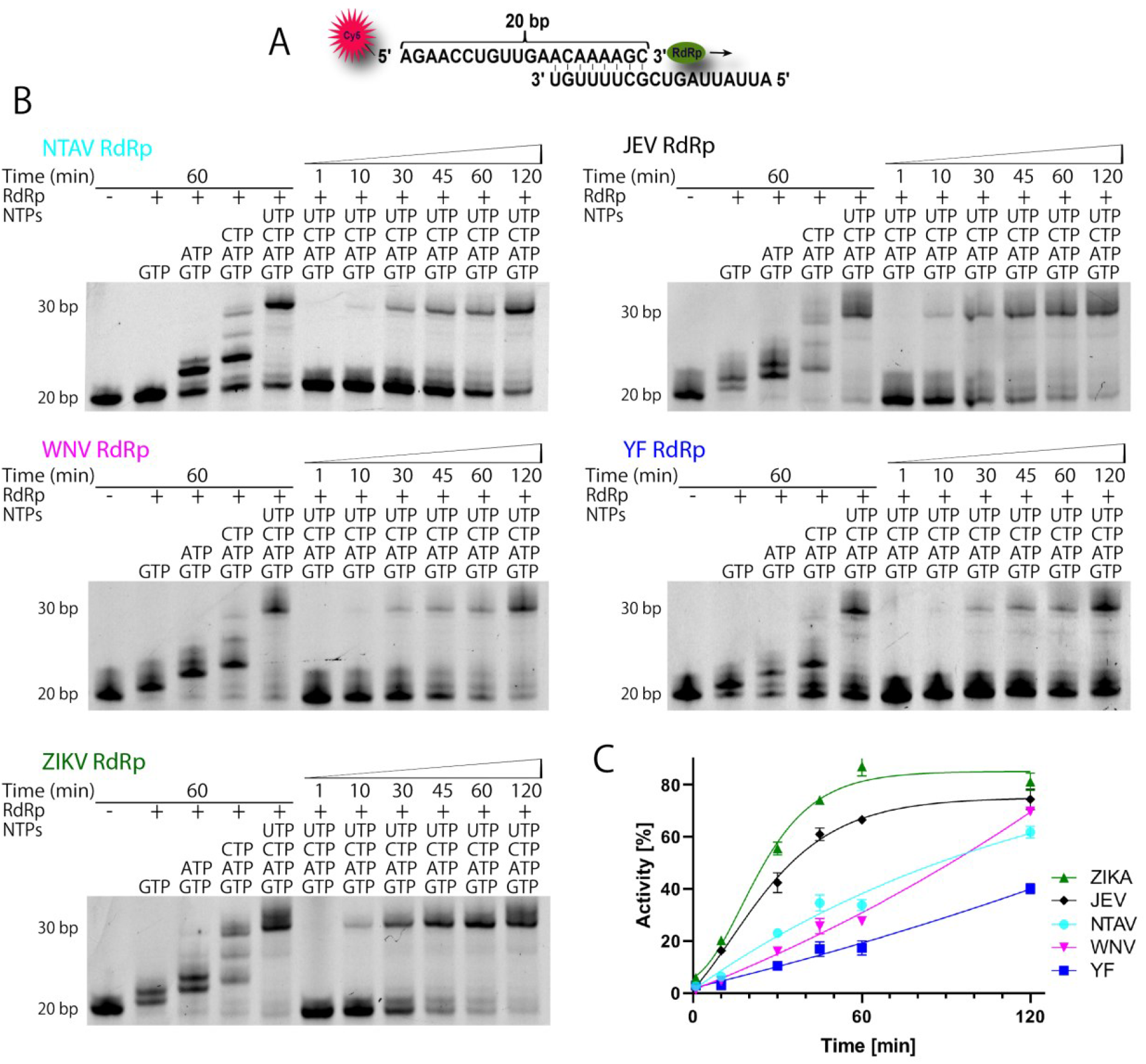
Analysis of polymerase activity of various flaviviral RdRps using a primer extension assay. A) RNA oligonucleotides used in this study. The fluorescent label (Cy5) at the 5’ end of one of the oligonucleotides is highlighted in red. The arrow indicates the direction of the primer extension. B) Incorporation of individual mixes of nucleotides in the RNA polymerase assay. The reaction contained 30 μM NS5 protein, 10 μM oligonucleotide duplex and was initiated by the addition of 10μM NTPs. All reactions were stopped at the given time-point and resolved on 20% denaturing PAGE gel. C) Graphical representation of RdRps activity (%) plotted against time (min).

### MTase enzymatic activities

We also aimed to compare enzymatic activities of the recombinant Ntaya MTase domain to those of better characterized flaviviruses (DENV3, WNV, ZIKV, TBEV, JEV and YFV). We prepared all these domains as recombinant proteins and measured their 2′-O-RNA MTase activity using ∼100 bp of their respective m7GpppA capped genomic RNA and SAM as substrates. For each methylated RNA molecule, one SAH molecule is produced and this SAH was quantified using mass spectroscopy. Surprisingly, we observed large differences among the various MTases. The most active was the Zika virus MTase, which converted 76% of substrate SAM to the product SAH in 85 minutes. NTAV, DENV, WNV, TBEV and JEV MTases showed 47% ± 3%, 45% ± 3%, 22% ± 2%, 9% ± 1% and 9% ± 1% of ZIKV MTase activity, respectively. Surprisingly, the activity of the YFV MTase was almost at the detection limit and almost inactive - only 3% ± 1% of ZIKV MTase activity (Figure 6).

**Figure 6:**
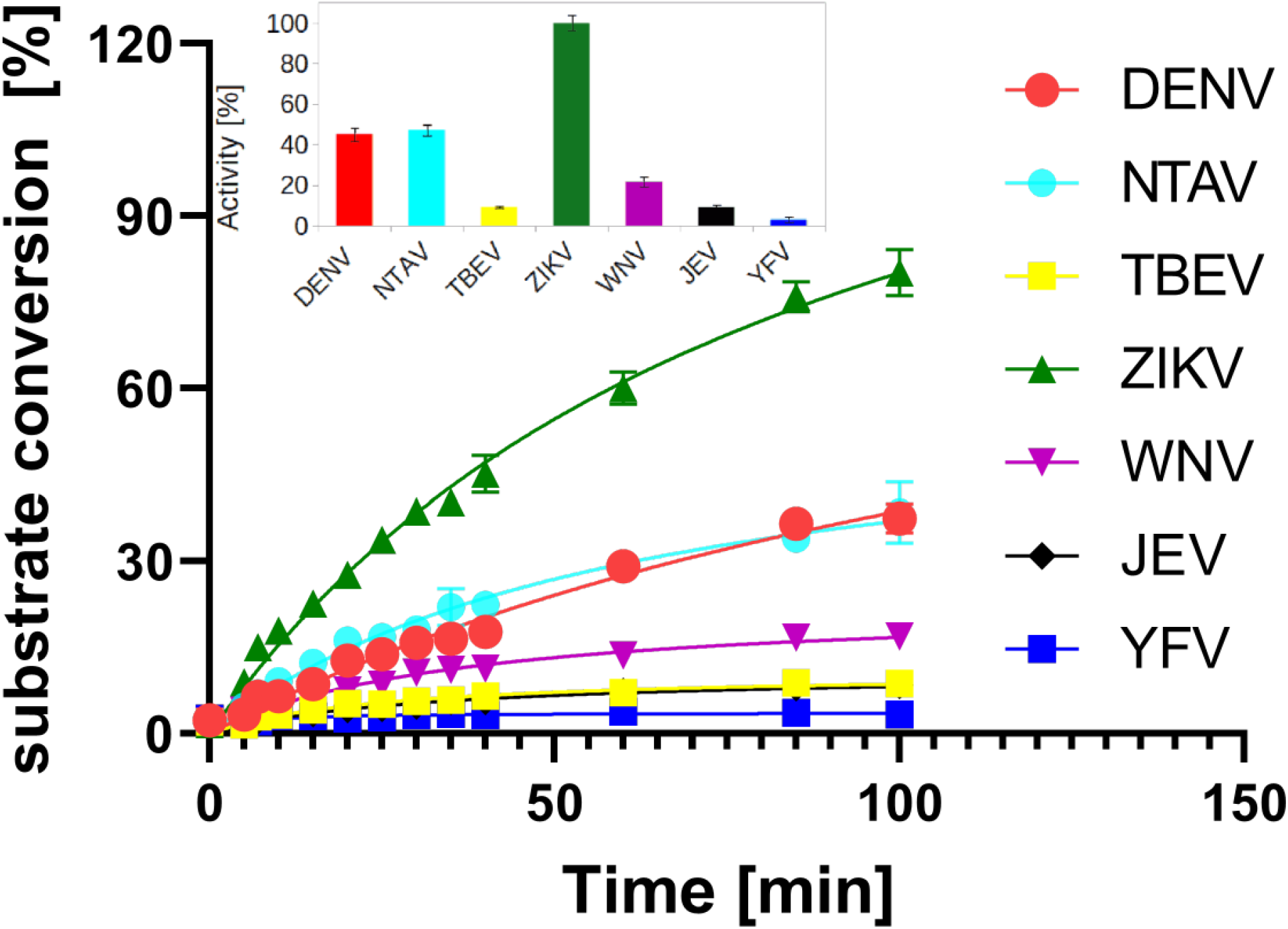
Analysis of the enzymatic activity of selected flaviviral MTases. The rate of MTase activity was measured as the amount of the substrate (SAM) converted to the product of the reaction (SAH). Data points are presented as mean values ± standard deviations from triplicates. For comparison of measured MTase activities the values of substrate conversion in 85 min reaction were expressed as percent of the value of ZIKV MTase substrate conversion.

## Discussion

Ntaya virus is primarily a zoonotic virus that is sometimes transmitted to humans and causes fever and headache [14]. It was discovered in the early fifties and is not considered too dangerous. In this respect, it resembles the Zika virus before the Zika epidemic that started in Brazil in 2015 [32]. Together with other recent outbreaks of +RNA zoonotic viruses (SARS, MERS, TBEV, SARS-CoV-2) and old foes such as YFV and WNV, there is a strong case advocating for considerable better understanding of +RNA viruses. In this study, we characterized the key protein responsible for RNA replication, NS5, of the Ntaya virus.

Our structural analysis revealed some differences in the conformations of several important regions such as the priming loop in the RdRp (Figure 2) or different conformations of Glu111 in the SAM binding pocket (Figure 3D). Glu111 forms a hydrogen bond with the 2′ hydroxyl group of the ribose ring in most of the structures of flaviviral MTases [33], but its ability to adopt a non-bonding conformation could help explain the mechanism of SAH leaving the active site because structurally, SAH and SAM are bound in the same way [16, 34-37]. Although there are differences among them, our structural comparison with previously available structures shows that the active sites in these + RNA viruses are conserved, indicating that there is significant evolutionary pressure to maintain these functional regions. This observation is encouraging because it suggests that a therapeutic compound active against one flaviviral enzyme should also be effective against all members of the flavivirus family.

Indeed, we have recently measured the activity of remdesivir triphosphate *in vitro* against various flaviviral polymerases, and it was very similar in the range of 0.3 - 2.1 µM [38]. Similar results were obtained for another more unusual inhibitor, PR673 [39]. These results correspond to our enzymatic analysis of recombinant flaviviral RdRps (Figure 5). The most active enzyme (ZIKV) was about 4x more active than the least active one (YFV). A somewhat different situation was observed among the MTase domains. Again, the MTase from ZIKV was the most active, but this time, more than an order of magnitude (30x actually) than the least active enzyme, which was again from YFV (Figure 6). These results are difficult to explain from the structural point of view. In any case, the RdRp has to synthesize the whole genome which is about 10 000 - 11 000 catalytic steps. At the same time, the MTase domain must perform one guanylyl transfer reaction, one N7 and one 2′-O methylation reactions. There might not be any evolutionary pressure for speed in the case of MTase domains explaining the differences we observe; a slow MTase domain could be just as good as a fast one.

## Concluding Remarks

RNA viruses, particularly +RNA viruses, pose a significant threat to humanity. To develop effective treatments against future epidemics, a thorough molecular understanding of these viruses is essential. Our study highlights the structural conservation of the enzymatic centers of both flaviviral RdRps and MTases, which offers promising opportunities for designing antivirals effective against all flaviviruses. Notably, although the enzymatic properties of recombinant MTases were diverse, all of the recombinant RdRps exhibited similar behavior.

## Material and methods

### Protein expression and purification

An artificial gene encoding the Ntaya NS5 protein (GeneBank: KF917539.1) was obtained from the European Virus Archive goes Global (EVAg). The sequence encoding the RdRp domain was cloned into pET28b vector using Gibson assembly. The resulting proteins contained an N-terminal 6× His-tag followed by TEV cleavage site. The sequence encoding the MTase domain was cloned into a home-made pSUMO vector [33] using restriction cloning (BamHI and XhoI sites). The resulting protein contained an N-terminal 8x-His-SUMO tag. All proteins were expressed and purified using our standard protocols for MTases and RdRps in E. coli [33, 40]. In brief, the genes were expressed in *E*.*coli* strain BL21-CodonPlus (DE3) RIL in LB medium supplemented with 50μM ZnSO_4_ and 1mM MgCl_2_. The bacteria were harvested by centrifugation, resuspended and sonicated in a lysis buffer (50mM Tris-HCl pH 8.0, 20mM imidazole, 500mM NaCl, 10% glycerol, 3mM β-mercaptoethanol). After lysis, the supernatant was immobilized on Ni-NTA agarose beads (Machery-Nagel), washed with lysis buffer supplemented with 1M NaCl and the protein was eluted using lysis buffer supplemented with 300mM imidazole.

For all the RdRps, the 6× His-tag was digested using TEV protease at 4°C overnight and the RdRps were further purified by affinity chromatography using HiTrap Heparin HP, HiTrap Q HP and Hi Trap SP HP columns (Cytiva). This was followed by size exclusion chromatography using Superdex 200 16/600 (GE Life Sciences) in 20 mM CHES pH 9.5, 800 mM NaCl, 10% (v/v) glycerol, 0.02% NaN3.

For the MTases, after elution from the Ni-NTA agarose beads, the proteins were supplemented with yeast sumo-protease Ulp1 and dialyzed against the lysis buffer overnight. The 8x-His-SUMO tag was removed by Ni-NTA agarose beads and the proteins were further purified by size exclusion chromatography using Superdex 75 16/600 (GE Life Sciences) in 25 mM HEPES pH 7.5, 500 mM NaCl, 5% glycerol and 1 mM TCEP. Finally, the pure proteins were concentrated to 4 mg/ml (RdRps) or 10 mg/ml (MTases) and stored at -80°C until needed.

### Crystallization

Crystals of Ntaya RdRp and MTase in complex with SAH grew in 7 days at 18°C in sitting drops consisting of 1:1 mixture (200 nl each) of the protein and the well solution (0.1 M Trizma/Bicine pH 8.5, 0.02M monosacharides, 10% (w/v) PEG4000, 20% (v/v) glycerol). GTP soaking was carried overnight in the presence of 1 mM Mg^2+^, the GTP concentration was 10 mM. The Ntaya MTase crystals in complex with sinefungin grew in two weeks in sitting drops prepared using the same procedure, but the well solution was 4.0 M sodium formate. These crystals did not require cryo-protection and were flash frozen in liquid nitrogen.

### Crystallographic analysis

The MTase datasets were collected using our home-source (rotating anode, Rigaku micromax-007 HF) while the RdRp dataset was collected at BESSY II electron storage ring operated by the Helmoltz-Zentrum Berlin (HZB) [41]. The data was integrated and scaled using XDS [42]. The structures of the NTAV MTase and NTAV RdRp were solved by molecular replacement using the structures of Zika MTase (pdb entry 5MRK) [33] and Yellow fever virus polymerase NS5A (pdb entry 6QSN) [22], respectively, as search models. The initial models were obtained with Phaser [43] from the Phenix package [44]. The models were further improved using automatic model refinement with Phenix.refine [45] followed by manual model building with Coot [46]. Statistics for data collection and processing, structure solution and refinement are summarized in SI Table 1. Structural figures were generated with the PyMOL Molecular Graphics System v2.0 (Schrödinger, LLC). The atomic coordinates and structural factors were deposited in the Protein Data Bank (https://www.rcsb.org).

### Primer extension polymerase activity assay

The polymerase activity of the NS5 RdRp domain was determined in a primer extension reaction using a fluorescently labeled primer (Cy5 5’-AGAACCUGUUGAACAAAAGC-3’) and a template (5’-AUUAUUAGCUGCUUUUGU-3’). The reaction was performed in a reaction mix containing 30 nM NS5 protein, 10 nM template/primer complex, 10 μM NTPs in the reaction buffer (5mM Tris-HCl pH 7.4, 10mM DTT, 0.5% Triton X-100, 1% glycerol, 3mM MnCl_2_) in a total volume of 20 µl. The data were quantified using ImageJ (NIH) and fitted to sigmoidal dose-response curves using. GraphPad Prism (Dotmatics).

### RNA preparation

The DNA templates (SI Table 2) for each flaviviral RNA were used for *in vitro* transcription in the presence of m7GpppA cap using the TranscriptAid T7 High Yield Transcription Kit (ThermoFisher Scientific). The obtained m7Gp3A capped RNAs were purified using RNA Clean and Concentrator (Zymo Research) and frozen in - 20°C until needed.

### MTase activity assay

The methyltransferase activity was measured using the MTase domains of NS5 proteins from NTAV, DENV3, WNV, ZIKV, TBEV, JEV and YFV. m7Gp3A capped RNA of the appropriate sequence for each virus (SI Table 2) was used as a substrate for the MTase assay. The reaction mixture contained 4 μM SAM and 4 μM m7Gp3A capped RNA in the reaction buffer (5 mM Tris pH 8.0, 1 mM TCEP, 0.1 mg/ml BSA, 0.005% Triton X-100, 1 mM MgCl_2_) and was started by the addition of the MTase to final concentration 0.5 μM in total volume 6 μl. The reaction mixture was incubated at 25°C for 0 – 100 min and analyzed using an Echo system coupled with a Sciex 6500 triple-quadrupole mass spectrometer operating with an electrospray ionization source. The rate of MTase activity was measured as the amount of the product of the reaction, SAH. The spectrometer was run in the multiple-reaction-monitoring (MRM) mode with the interface heated to 350 °C. The declustering potential was 20 V, the entrance potential was 10 V, and the collision energy 28 eV. 10 nl of each sample was injected into the mobile phase (flow rate of 0.40 ml/min; 100% methanol). The characteristic product ion of SAH (m/z 385.1 > 134.1) was used for quantification.

## Acknowledgment

We thank the Helmholtz-Zentrum Berlin für Materialien und Energie for the allocation of synchrotron radiation beamtime. This research was funded by the Czech Science Foundation [21-25280S]; the project the National Institute Virology and Bacteriology (Programme EXCELES, Project No. LX22NPO5103) - Funded by the European Union - Next Generation EU, and by the Grant Agency of Charles University (Grant No. 408422). The Academy of Sciences of the Czech Republic, RVO: 61388963, is also acknowledged.

## Conflict of interests

The authors declare no conflict of interests.

## Notes

### Competing Interest Statement

The authors have declared no competing interest.

