## Supplementary Information for "Structural and functional insights in flavivirus NS5 proteins gained by the structure of Ntaya virus polymerase and methyltransferase"

| Crystal | MTase + sinefungin | MTase + SAH + GTP | RdRp |
|---|---|---|---|
| PDB accession code | 8BXK | 8CQH | 7ZIU |
| **Data collection and processing** | | | |
| Space group | $P2_12_12_1$ | $P2_1$ | $P2_1$ |
| Cell dimensions - a, b, c (Å) | 55.1, 110.1, 116.2 | 38.3, 71.7, 50.4 | 64.3, 79.8, 133.3 |
| Cell dimensions - α, β, γ (°) | 90, 90, 90 | 90, 93.2, 90 | 90, 93.3, 90 |
| Resolution range (Å) | 39.96 - 2.31 (2.39 - 2.31) | 27.40 - 2.00 (2.07 - 2.00) | 47.58 - 2.80 (2.90 - 2.80) |
| No. of unique reflections | 31652 (1077) | 18427 (1817) | 33010 (3220) |
| Completeness (%) | 87.16 (34.21) | 99.86 (100.00) | 98.80 (97.66) |
| Multiplicity | 6.3 (6.4) | 13.3 (12.9) | 6.9 (6.8) |
| Mean I/σ(I) | 16.4 (3.7) | 22.1 (7.2) | 6.13 (0.94) |
| $CC_{1/2}$ | 0.998 (0.966) | 0.998 (0.964) | 0.983 (0.529) |
| CC* | 1 (0.991) | 1 (0.991) | 0.996 (0.832) |
| **Structure solution and refinement** | | | |
| R-work (%) | 20.03 (20.50) | 15.22 (15.16) | 23.74 (36.83) |
| R-free (%) | 23.46 (23.50) | 18.98 (20.58) | 28.88 (45.41) |
| R.m.s.d. - bonds (Å) / angles (°) | 0.003 / 0.46 | 0.006 / 0.87 | 0.002/0.43 |
| Average B factors ($Å^2$) | 28.56 | 14.77 | 57.33 |
| Clashscore | 11.61 | 3.65 | 4.28 |
| Ramachandran favored/outliers (%) | 98.6 / 0 | 99.2 / 0 | 98.1 / 0 |

SI Table 1. Statistics of crystallographic data collection and refinement. Numbers in parentheses refer to the highest resolution shell.

| Flavivirus | Template for *in vitro* transcription |
|---|---|
| DENV3 | CAGTAATACGACTCACTATAGttgttagtctacgtggaccgacaagaacagtttcgactcggaagcttgcttaa<br>cgtagtgctgacagtttttattagagagcagatctctga |
| NTAV | CAGTAATACGACTCACTATAGaagttcatctgtgtgaacttcgtgattgacagctcaacacgagtgcgggcaa<br>ccgtaaacacagtttgaacgtttttggagagagactact |
| TBEV | CAGTAATACGACTCACTATAGattttcttgcacgtgcgtgcgtttgcttcggacagcattagcagcggttggtttg<br>aaagaaatattcttttgtttttaccagtcgtgaacgtgttgagaaaaagacagcttaggagaacaagagctgggg |
| ZIKA | CAGTAATACGACTCACTATAGttgttgatctgtgtgagtcagactgcgacagttcgagtctgaagcgagagcta<br>acaacagtatcaacaggtttaatttggatttggaaacgagagtttctggtc |
| WNV | CAGTAATACGACTCACTATAGtagttcgcctgtgtgagctgacaaacttagtagtgtttgtgaggattaacaaca<br>atta acacagtgcgagctgtttcttggcacgaagatctcg |
| JEV | CAGTAATACGACTCACTATAGaagtttatctgtgtgaacttcttggcttagtatcgttgagaagaatcgagagatt<br>agtgcagtttaaacagtttttagaacggaagataacc |
| YFV | CAGTAATACGACTCACTATAGtaaatcctgtgtgctaattgaggtgcattggtctgcaaatcgagttgctaggca<br>ataaacacatttggattaattttaatcgttcgttgagcgattagcagagaactgaccagaac |

SI Table 2. DNA sequences used as templates for *in vitro* transcription. Sequence of T7 promotor is in capitals, first base in the transcript is the G (underlined).

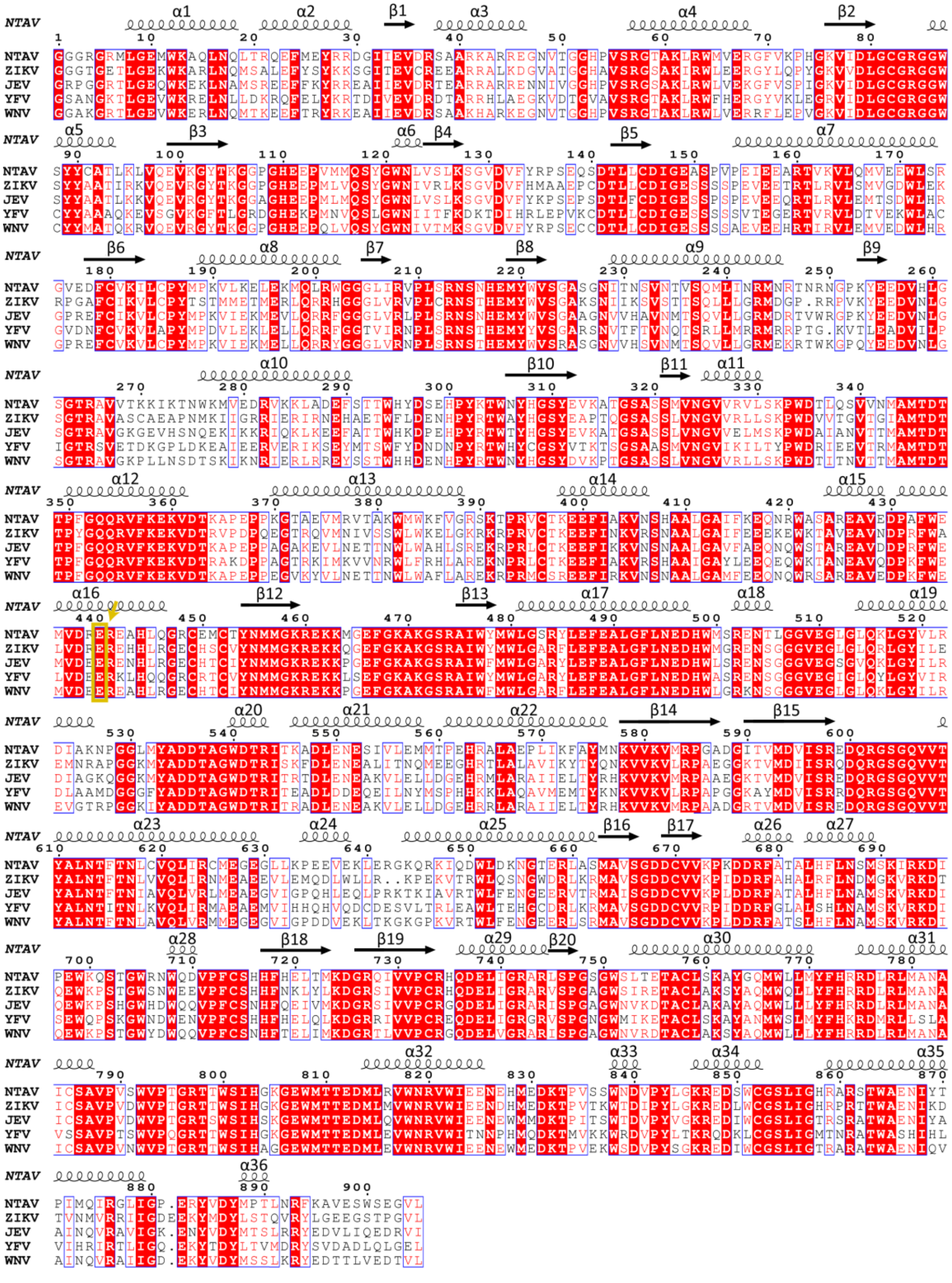

SI Figure 1: **Primary sequence alignment of flaviviral NS5s.** Conserved residues are highlighted in red. Glu440 is highlighted by a yellow arrow. Secondary structure elements of Ntaya NS5 are shown above the sequence. The alignment was made in ESPript 3.0 online program (https://espript.ibcp.fr/ESPript/ESPript/).

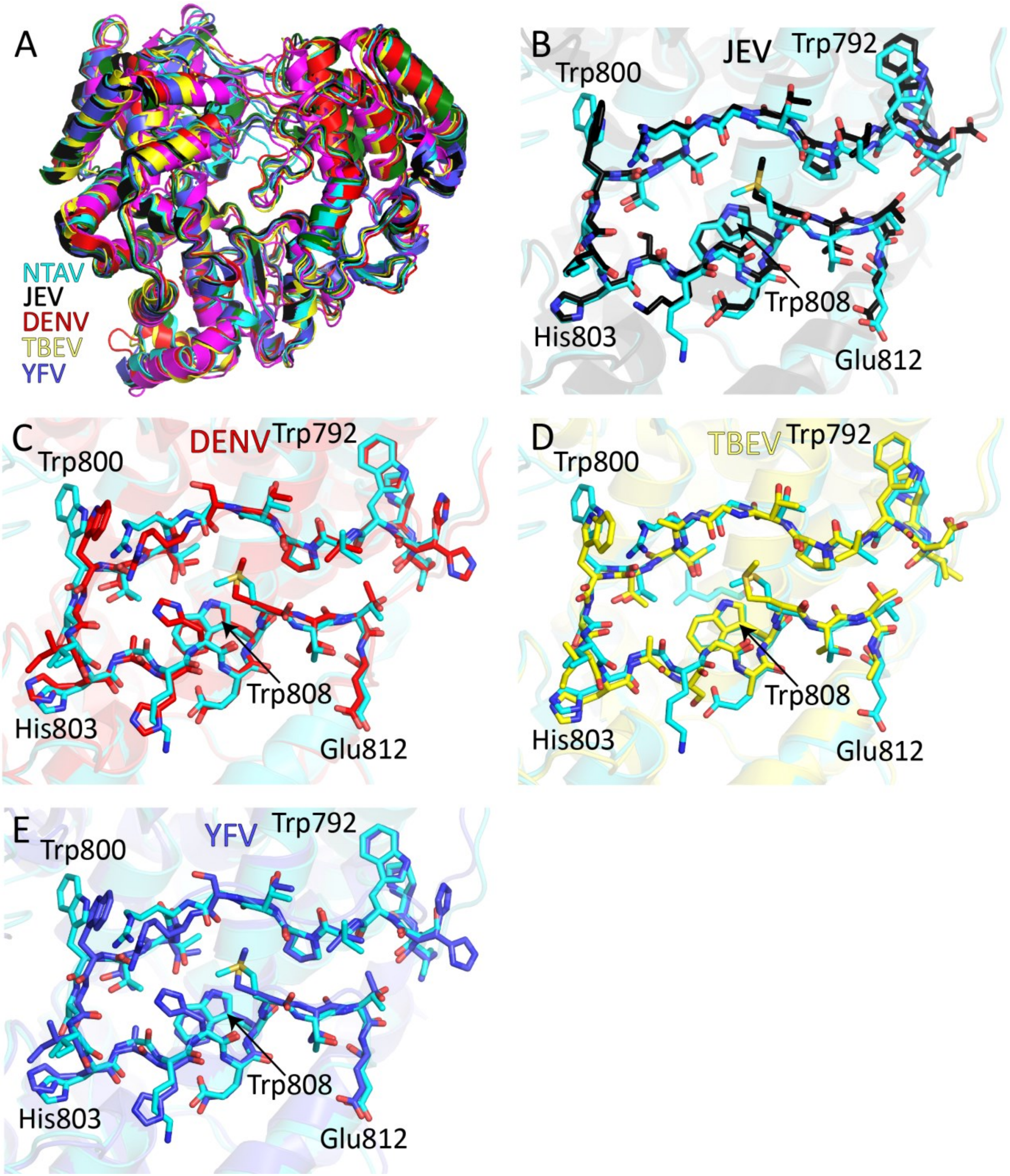


SI Figure 2: **Structural alignment of Flaviviral RdRp domains.** A) The overall structural alignment of Ntaya (cyan), Japanese encephalitis (black, PDB: 4K6M), Dengue 3 (red, PDB: 5CCV), Tick-borne encephalitis (yellow, PDB:7D6M) and Yellow fever (blue, PDB: 6QSN) RdRps. B-E) Structural superposition of priming loops of flaviviruses mentioned above.

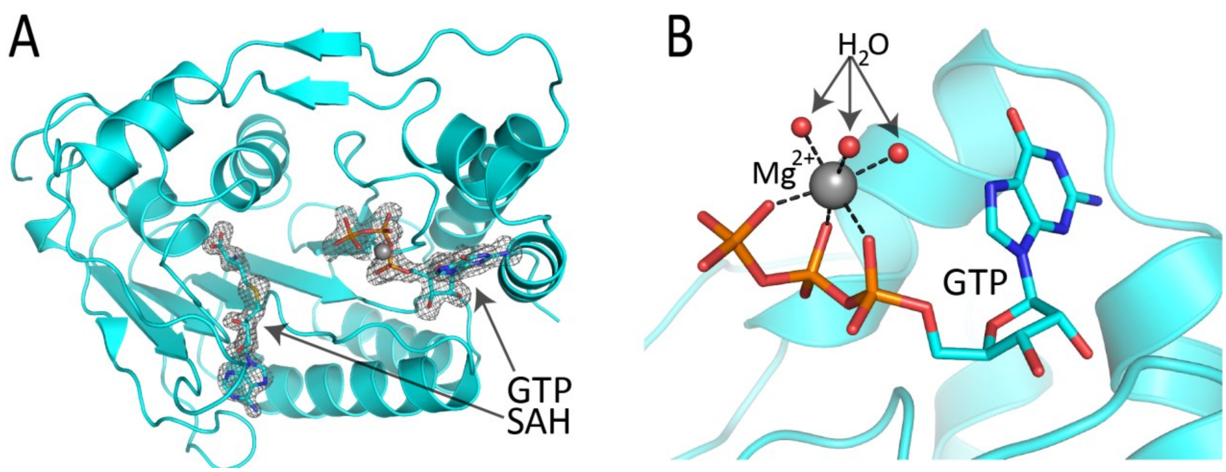


SI Figure 3: A) Overall structure of Ntaya MTase domain with labeled secondary structure elements and Fo-Fc omit map of electron density at 3σ with GTP and SAH excluded from the calculation. B) Detailed view of coordination of the magnesium ion by three water molecules and three phosphate groups of the GTP.
